# Brain-synthesized estrogens regulate cortical migration in a sexually divergent manner

**DOI:** 10.1101/654731

**Authors:** Katherine J. Sellers, Matthew C.S. Denley, Atsushi Saito, Atsushi Kamiya, Deepak P. Srivastava

## Abstract

Estrogens play an important role in the sexual dimorphisms that occur during brain development, including the neural circuitry that underlies sex-typical and socio-aggressive behaviors. Aromatase, the enzyme responsible for the conversion of androgens to estrogens, is expressed at high levels during early development in both male and female cortices, suggesting a role for brain-synthesized estrogens during corticogenesis. This study investigated how the local synthesis of estrogens affects neurodevelopment of the cerebral cortex, and how this differs in males and females by knockdown expression of the *Cyp19a1* gene, which encodes aromatase, between embryonic day 14.5 and postnatal day 0 (P0). The effects of *Cyp19a1* knockdown on neural migration was then assessed. Aromatase was expressed in the developing cortex of both sexes, but at significantly higher levels in male than female mice. Under basal conditions, no obvious differences in cortical migration between male and female mice were observed. However, knockdown of *Cyp19a1* increased the number GFP-positive cells in the cortical plate, with a concurrent decrease in the subventricular zone/ventricular zone in P0 male mice. The opposite effect was observed in females, with a significantly reduced number of GFP-positive cells migrating to the cortical plate. These findings have important implications for our understanding of the role of fetal steroids for neuronal migration during cerebral cortex development. Moreover, these data indicate that brain-synthesized estrogens regulate radial migration through distinct mechanisms in males and females.

## Introduction

The unique organization and architecture of the cerebral cortex is established during embryonic development. This organization is achieved in an “in-side-out” fashion, with inner, or deep layers forming first, and outer, or superficial layers developing last (Evsyukova *et al.*, 2013). The development of the cortex and its laminar organization is controlled in large part by the coordinated processes of neurogenesis and cell migration. The process of radial neuronal migration is a multi-step process (Evsyukova *et al.*, 2013). Initially, newly born neurons, generated from neural stem cells, detach from the apical surface of the germinal ventricular zone (VZ). These neurons adopt a multi-polar morphology and move into the intermediate zone (IZ) (Noctor *et al.*, 2004). Here, neurons develop a bi-polar shape, and migrate along radial glia to their final position in the cortical plate (CP) (Kawauchi, 2015). Once in this position, neurons can begin to form synaptic connections, and thus contribute to circuit formation (Evsyukova *et al.*, 2013).

Several areas of the brain undergo sexually dimorphic development. During adolescent development, the brain undergoes differential development trajectories that lead to sexual dimorphism in total cerebral volume and different local grey matter nuclei volumes (Kaczkurkin *et al.*, 2019). Notably, female total cerebral volume peaks earlier in adolescence than males (Lenroot *et al.*, 2007). In addition, analysis of neural circuitry and behavior reveals that different systems and regions within the brain are engaged in a sex-dependent manner and specific nuclei within the brain are thought to be responsible for sex-specific behaviors (Gillies & McArthur, 2010; Choleris *et al.*, 2018). The hippocampus and basolateral nucleus of the amygdala are necessary for sex-dependent learning (Bangasser & Shors, 2007; Waddell *et al.*, 2008). However, other regions of the brain also display sexually dimorphic neural circuitry, including the bed nucleus of the stria terminalis (thalamus) and the medial prefrontal cortex (frontal lobe) (Bangasser & Shors, 2007; Maeng *et al.*, 2010). Structural differences at a macroscopic level are witnessed. For example, sexual dimorphism has been recorded in the preoptic area (hypothalamus), uncinate nucleus (hypothalamus), olfactory bulb (rostral frontal lobe), anterior commissure (white matter tract connecting hemispheres), interthalamic adhesion (thalamus), and mammillary body (diencephalon) (Allen & Gorski, 1991; Garcia-Falgueras & Swaab, 2008; Savic *et al.*, 2010; Oliveira-Pinto *et al.*, 2014).

Estrogens, in particular 17β-estradiol (estradiol), are integral in establishing sexual dimorphisms during brain development, including the neural circuitry that underlies sex-typical and socio-aggressive behaviors (Gillies & McArthur, 2010; McCarthy *et al.*, 2018). Sex-hormones, especially estrogens such as estradiol, have consistently shown cognitive-enhancing properties and morphological regulatory responsibilities (Gillies & McArthur, 2010; Sellers *et al.*, 2015a; Choleris *et al.*, 2018). By modulating spinogenesis, synaptogenesis, and synaptic connectivity, estrogen enacts rapid changes to the neural circuitry (Saldanha *et al.*, 2011; Srivastava *et al.*, 2013; Sellers *et al.*, 2015a; Sellers *et al.*, 2015b). Steroid-dependent plasticity and circulating steroid hormones impact neuronal volume and numbers (Balthazart & Ball, 2006; Saldanha *et al.*, 2011; Srivastava *et al.*, 2013). Interestingly, estrogens also stimulate the proliferation and differentiation of neural progenitors and neuronal populations (Brannvall *et al.*, 2002; Okada *et al.*, 2010; Denley *et al.*, 2018). Furthermore, estradiol has recently been shown to regulate neurite outgrowth in immature cortical neurons derived from human induced pluripotent stem cells (Shum *et al.*, 2015). Importantly, estrogen receptor-beta (ERβ) knockout animals have been reported to display abnormal neuronal migration in the neocortex (Wang *et al.*, 2003). However, whether the regulation of migration via ERβ is due to systemic or brain-synthesised estradiol is unclear. Similarly, whether estradiol influences neuronal migration in male and females is not known. Nevertheless, these lines of evidence indicate that estrogens play a role in during the development of the cortex.

Aromatase, the enzyme responsible for the conversion of androgens and cholesterol to estrogens, is expressed across species and is highly expressed during early development in both male and female cortices, suggesting a role for brain-synthesized estrogens during corticogenesis. Aromatase is believed to be the source of *de novo* estradiol synthesis in areas such as the hippocampus and cortex (MacLusky *et al.*, 1994; Saldanha *et al.*, 2011; Srivastava *et al.*, 2013; Lu *et al.*, 2019). We previously demonstrated that factors involved in estrogenic signalling, including aromatase, are present during the development of the human cortex (Denley *et al.*, 2018). In addition, aromatase-mediated estradiol signalling is required in fear-learning regulated by the basolateral amygdala and leads to sexually dimorphic plastic responses (Bender *et al.*, 2017). In the developing mammalian brain, supporting cells such as astrocytes and radial glial cells also express aromatase (Martinez-Cerdeno *et al.*, 2006; Yague *et al.*, 2006; Yague *et al.*, 2008). Studies in zebrafish demonstrated that radial glial cells expressing aromatase divide into neurons and that neural stem cells in the ventricular layer also express aromatase (Pellegrini *et al.*, 2007). Taken together, these data provide strong evidence for a role for aromatase in differentiation and neurogenesis.

The purpose of this study was to investigate how the local synthesis of estrogens affects neurodevelopment of the cerebral cortex, and specifically the somatosensory cortex, and how this differs in males and females. Using an shRNA-approach, we knocked-down the *Cyp19a1* gene, which encodes aromatase, from an early developmental stage and assessed the loss of aromatase, and thus the ability to locally synthesise estrogens had on neural migration. Analysis of neural migration revealed a sex specific effect of aromatase loss on neural migration. Taken together, these data contribute to the current evidence that brain-synthesised estrogens play a role in the development of the cortex, and moreover, add to the growing appreciation of sexually dimorphic niches in the brain.

## Materials and Methods

### Plasmid construction and shRNA validation

Four shRNAs against Mus musculus *Cyp19a1* and one control scrambled shRNA were obtained from Origene (Rockville; Cat No. TG509276). These plasmids express both shRNA under the control of the U6 promoter and turboGFP under the control of a CMV promoter. A myc-DDK-tagged aromatase construct (pCMV6-myc-DDK-aromatase) was purchased from Origene (Rockville; Cat. No. MR224509); the pCAG-eGFP has previously been described (Srivastava *et al.*, 2012b). The effectiveness of each shRNA was validated by testing the ability of each construct to knockdown myc-aromatase expression in hEK293T cells. Briefly, hEK293 cells were grown to 40% confluency before transfection of myc-aromatase with or without shRNA constructs using Lipofectamine 2000 (Lifetechnologies, UK). Transfections were allowed to procced for 48 hours, after which cells were lysed and prepared for western blotting as previously described (Srivastava *et al.*, 2012a).

### In utero electroporation

*In utero* electroporation targeting the somatosensory cortex was preformed according to previously published protocol (Niwa *et al.*, 2010; Saito *et al.*, 2016). All experiments were performed in accordance with the institutional guidelines for animal experiments. Embryos were electroporated at E14.5. Pregnant CD1 mice (obtained from Charles River) were anesthetized by intraperitoneal administration of a mixed solution of Ketamine HCl (100 mg/kg), Xylazine HCl (7.5 mg/kg), and Buprenorphine HCl (0.05 mg/kg). The uterine sacs were exposed by laparotomy. In each pregnant animal, aromatase-shRNA, scramble-shRNA (1 μg/μl) or eGFP (0.5 μg) were injected into the left ventricle of the embryo with a glass micropipette made from a microcapillary tube (GD-1; Narishige). Control embryos were injected eGFP (1.0 μg/μl) into the right ventricle in the same manner. The head of the embryo was held between the electrodes (Nepagene) placed over the posterior forebrain with the positive electrode positioned above sight of injection. Electrode pulses (33V; 50 ms) were charged four times at intervals of 950 ms with an electroporator (CUY21EDIT; Nepagene). After electroporation the uterine horn was replaced in the abdominal cavity to allow the embryos to continue to develop. A total of 6-15 embryos were electroporated in each of the 5 pregnant animals. Brains were extracted from P0 pups and assessed for GFP expression in correct location prior to fixation. Brains were fixed by overnight immersion in 4% paraformaldehyde in 0.1 M phosphate buffer saline (PBS), and then cryoprotected/stored at 4°C in 30% sucrose in PBS.

### Genotyping

Genomic DNA extracted from tail biopsies of postnatal day 0 mice (0.5 cm of tail removed from mice under anaesthesia) was analyzed using polymerase chain reaction. The tissue was digested overnight at 55°C in lysis buffer containing 100 mM Tris-HCl pH 8.5, 5 mM EDTA, 0.2% SDS, 200 mM NaCl, and 0.1 mg/ml proteinase K (Roche, Basel, Switzerland). Undigested material was removed by centrifugation. The pellet was re-suspended in nuclease-free water and the absorbance was measured using a NanoDrop ND1000 spectrophotometer (Nanodrop Technologies, Wilmington, DE, USA). Sex genotyping was performed using primers for Sry sex determining region of the Y-chromosome. Primers as follows: forward SRY Forward: TTG TCT AGA GAG CAT GGA GGG CCA TGT CAA, and reverse SRY Reverse: CCA CTC CTC TGT GAC ACT TTA GCC CTC CGA. These identify a 273 base pair PCR product. The PCR reagents as follows: 2.5 µl 10× PCR buffer (Amersham Pharmacia Biotech); 0.2 mM dNTP mix; 0.25 µM forward and reverse primer set 1; 1 µM forward and reverse primer set 2; 0.625 U Taq DNA polymerase; and 2 µl DNA template (or 2 µl sterile PCR-grade H_2_O for PCR control sample); the volume was made up to 25 µl using sterile PCR-grade H_2_O. PCR was performed using a Peltier Thermal Cycler (PTC-200, MJ Research Inc., Watertown, MA) The resulting amplicons were resolved on 1.5% agarose gels and visualized using ethidium bromide staining with a GelDoc transilluminator (BioRad).

### Immunohistochemistry (IHC)

Brains were mounted in OCT embedding media (Bright) and cut into 14 µm sections across the coronal plane using a cryostat (Leica CM 1860 UV, Ag Protect) and collected on SuperFrost Plus microscope slides (Thermo Scientific). Immunohistochemistry (IHC) was carried out as previously described (Jones *et al.*, 2019). In brief, sections were simultaneously permeabilised and blocked in 0.1% Triton-X with 2% normal goat serum in PBS for 1 h at room temperature, in a humidified chamber. They were then incubated overnight at 4°C in a humidified chamber with primary antibodies against chicken GFP (1:1000; Abcam #ab13970) and rabbit aromatase (1:100 Abcam #ab18995). Sections were then counterstained the appropriate secondary antibodies and counter stained with DAPI (ThermoFischer D1306). Images were captured and analyzed as described below.

### Image acquisition and data analysis

Confocal images of IHC stained P0 sections was carried out using a Nikon Spinning Disk confocal equipped with either a 20X or 40X objective. Image z-stacks were acquired at z-intervals optimised for the specific objective. Images to be used for subsequent intensity-based analysis were acquired using identical acquisition parameters. For migration analysis, cortical sub-sections were identified by DAPI staining and regions of interest (ROIs) were identified in the 488 nm channel using the epifluorescence microscope. The effect of knockdown of aromatase on migration was analyzed by quantitative bin analysis according to previously published methods (Kubo *et al.*, 2010). In brief, the cortex was divided into ten equal sections (bins) and percentage of GFP^+^ cells within each bin was determined. For each condition, a minimum of three images were collected over at least six section.

Analysis of aromatase expression was carried out on sections immunostained for aromatase and DAPI (endogenous aromatase expression) or immunostained for GFP, aromatase and DAPI (assessment of aromatase knockdown). ROIs were determined either as described above or limited to GFP^+^ cells. Images were background subtracted, and the mean intensity of aromatase staining determined for five intendent ROIs per image; three ROIs of background staining were also measured for each section. The mean intensity for each section was normalised to background staining (average of 3x background ROIs + 2x StDev). Between 3-4 sections per brain; 3 brains per condition was used for these analyses.

## Results

### Sex differences in aromatase expression but not in migration during cortex development

During human embryonic neocortical development, aromatase expression is transient but typically peaks during the last trimester and early postnatal period and then drops dramatically following embryonic maturity (Montelli *et al.*, 2012; Denley *et al.*, 2018). Similarly, estradiol has been measured in pre- and post-natal cortex of male and female rats (Konkle & McCarthy, 2011). Consistent with these findings, we found that aromatase was expressed in the developing cortex of mice (**Figure 1A**). In order to establish whether there was a sex difference in the expression of aromatase in the developing cortex, we examined expression of this enzyme in male and female P0 mice. Sex was established by the assessment of sex-determining region Y (*sry*) gene expression (**Figure 1B**). IHC analysis revealed aromatase positive cells in the SVZ/VZ, IZ and CP of male and female P0 mice (**Figure 1C**). Aromatase could be observed in the cell body (red arrows) as well as in dendrite-like processes (yellow arrow heads) of cells within the CP in both male and female P0 mice (**Figure 1D**). In line with previous studies (Montelli *et al.*, 2012), quantification of aromatase expression revealed that aromatase expression was higher in the developing cortex of male compared to female P0 mice (**Figure 1E**). Taken together, these data indicate that aromatase is expressed in the developing cortex of male and female mice, suggesting a potential role for brain-synthesised estrogens during this developmental time point.

**Figure 1.**
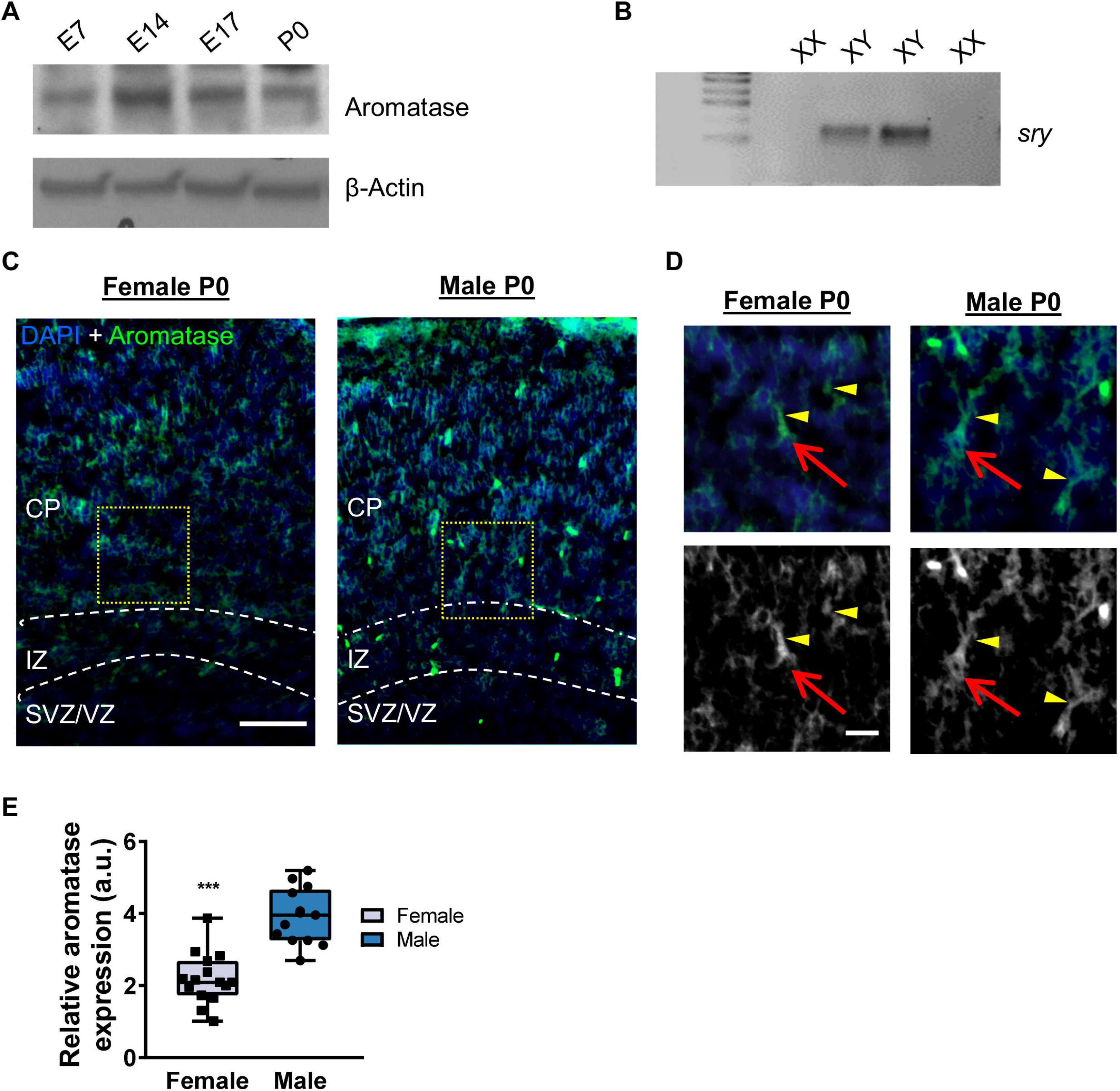
Aromatase is differentially expressed in the developing female and male cortex. **(A)** Western blot of lysates taken from developing cortex demonstrating aromatase expression at different stages of development. **(B)** Representative RT-PCR analysis for *sry* gene used to determine sex of P0 animals used in subsequent analysis. **(C)** Representative confocal image of aromatase staining in P0 cortex of female and male. Aromatase positive cells can be observed in the SVZ/VZ, IZ and CP in both female and male developing cortex. Yellow dotted boxes indicate high magnification images shown in D. **(D)** High magnification images from C. Red arrows indicate aromatase positive immunoreactivity within cell body of developing neurons in either female or male P0 cortex. Yellow arrow heads indicate aromatase expression within processes of developing neurons in P0 cortex of both sexes. **(E)** Quantification of relative aromatase expression throughout P0 cortex of female and male. This revealed that males have a significantly higher expression of aromatase at this stage of development within the cortex. Conditions were compared by Student’s t test; **P<0.001; n=13-15 sections from 8 pups/sex. Data are presented as Box and Whisker plots with error bars showing minimum and maximum data points; each data point represents an individual section. Scale bars = 100 μm (C) and 20 μm (D).

As our data indicated that aromatase is more highly expressed in males compared to females, we reasoned that this may be reflected in a difference in neuronal migration between sex. Therefore, to analyze the effect of sex on neuronal migration, we *in utero* electroporated embryos at E14.5 with a GFP expressing plasmid (pCAG-eGFP), and quantified the positions of GFP-positive (GFP+) cells in P0 brains as previously described (Kubo *et al.*, 2010). Sex were again determined by assessing expression of *sry.* Analysis of neuronal migration revealed that the differential expression of aromatase did not confer a difference in the migration of GFP+ cells in either sex under these control conditions (**Figure 2A**). This observation is confirmed by the relatively parity of GFP+ cells throughout the developing P0 cortex (**Figure 2B**). These data indicate that there are no differences in neuronal migration between male and female mice at P0.

**Figure 2.**
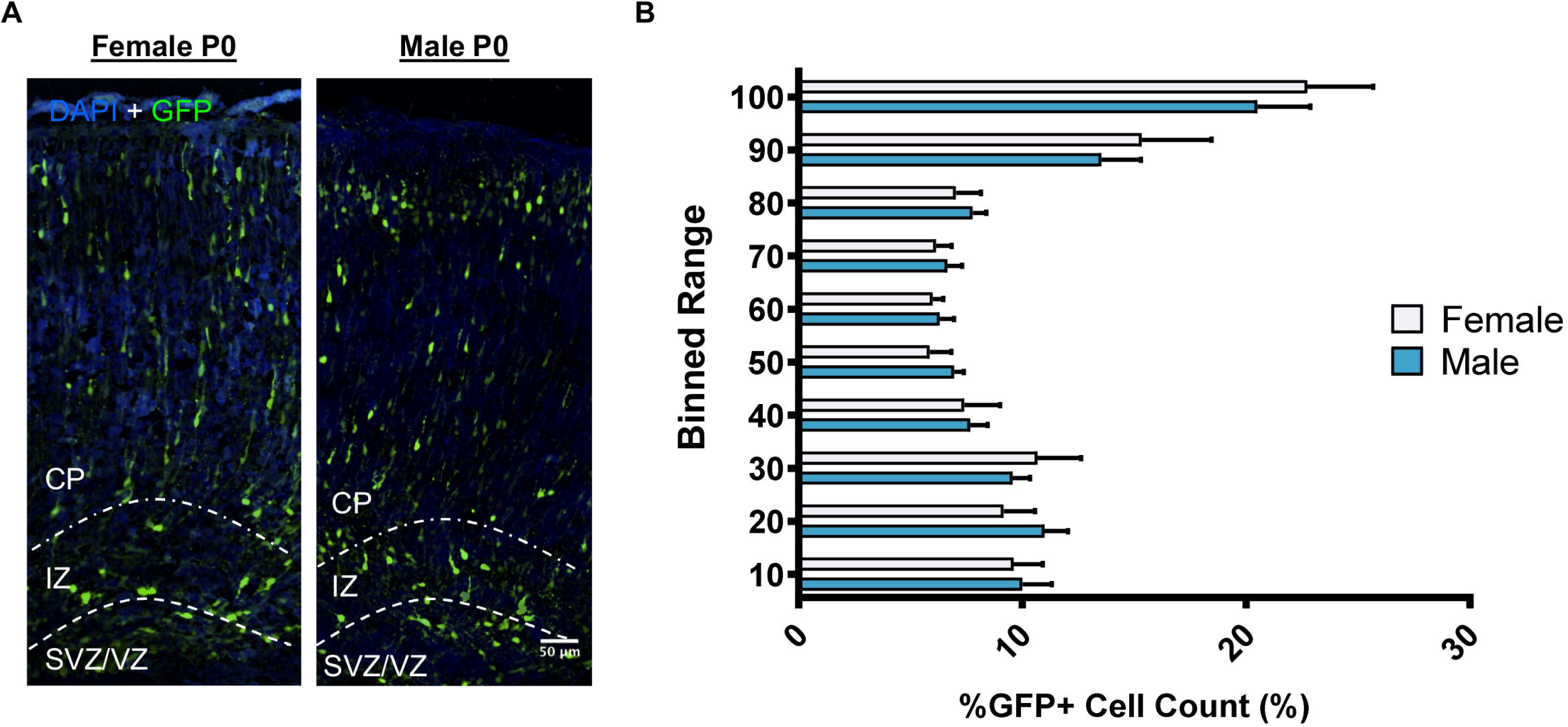
Neuronal migration does not differ between female and male in the developing cortex. **(A)** Representative confocal images of P0 cortex of female and males demonstrating the distribution of GFP+ cells. **(B)** Quantification of GFP+ cells throughout the developing cortex. This revealed that there was a similar distribution of GFP+ positive cells the P0 cortex of female and male mice. Scale bar = 50 μm.

### Validation of aromatase knockdown

Although our data indicates that there is no difference in neuronal migration between male and females under control conditions, previous studies using ERβ knockout animals have suggested a role for estradiol in cortical development (Wang *et al.*, 2003). Furthermore, estradiol has been shown to regulate proliferation and differentiation of neural progenitor cells (Denley *et al.*, 2018). Therefore, we were interested in understanding whether brain-synthesised estrogens may contribute to neuronal migration in either male or female mice. In order to do this, we employed a short hairpin RNA interference (shRNA) approach to selectively knockdown expression of *Cyp19a1*, the gene encoding aromatase. The efficiency of four individual shRNA to knockdown aromatase was first established in hEK203 cells. Myc-tagged aromatase was exogenous expressed in hEK293 cells in the presence of shRNA for aromatase or a control (scramble) shRNA (**Figure 3A**). Of the four shRNA tested shRNA_c (herein referred to as shRNA_arom) reduced myc-aromatase expression by ∼60% (**Figure 3A and B**) and was used in subsequent experiments.

**Figure 3.**
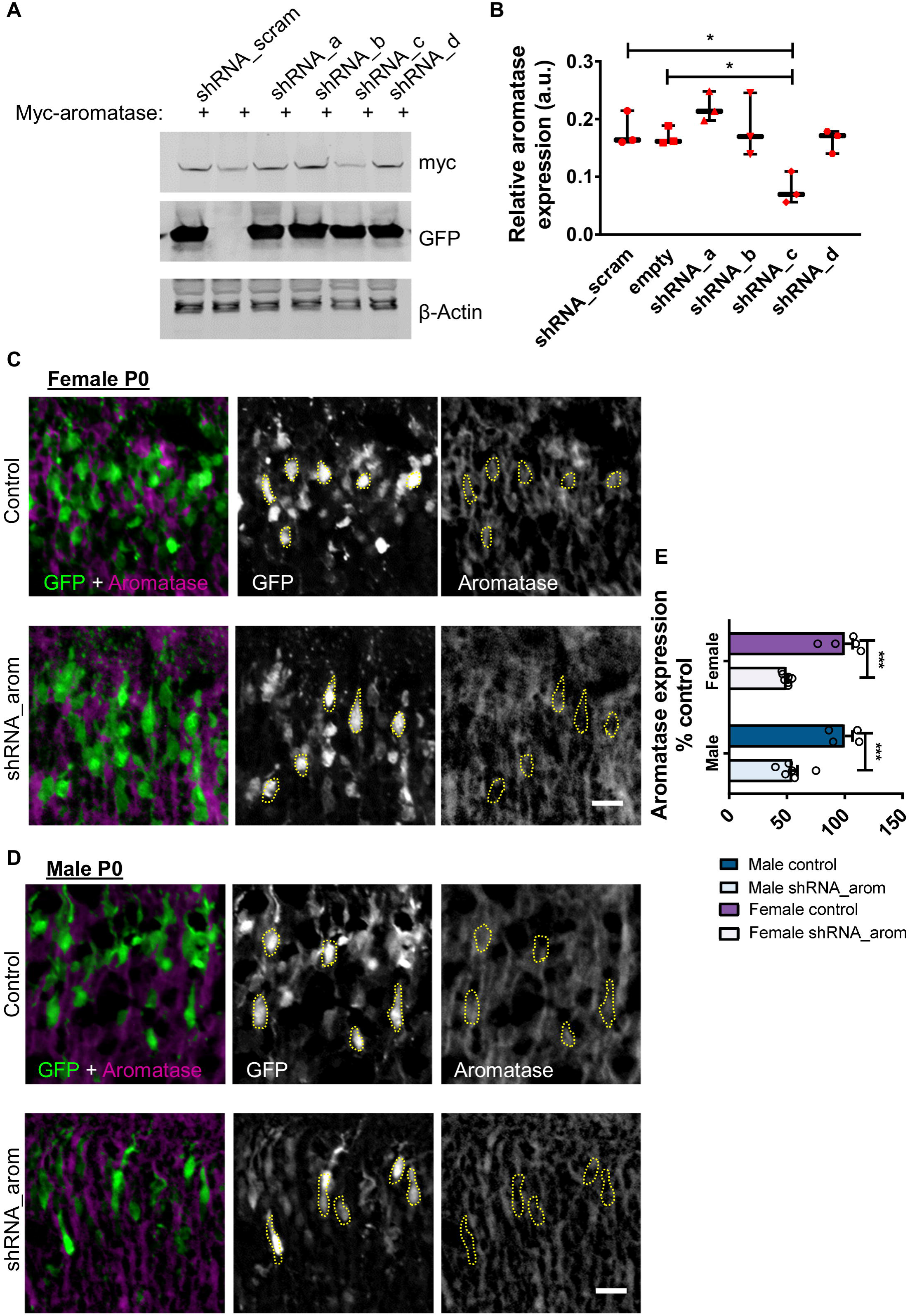
Optimization of *Cyp19a1* knockdown *in vivo*. **(A)** Western blot of cell lysates taken from hEK293 cells expressing myc-aromatase in the presence or absence of shRNA against *Cyp19a1* (shRNAa-d) or a control construct (shRNA_scram). **(B)** Quantification of myc-aromatase expression reveals that shRNA_c (aka shRNA_arom) produced ∼60% knockdown of exogenous aromatase. Conditions were compared by one-way ANOVA; F(5,12)=7.077, p=0.003, Tukey Post Hoc, *, p < 0.05; n = 3 independent cultures. Data are presented as Box and Whisker plots with error bars showing minimum and maximum data points; each data point represents an individual experiment. **(C and D)** Representative confocal images of female (C) or male (D) P0 cortex *in utero* electroporated at E14.5 with either control (shRNA_scram) or shRNA_arom and co-stained for aromatase and GFP. GFP was used to identify cells expressing control or *Cyp19a1* shRNA. **(E)** Quantification of aromatae expression in GFP+ cells in control or shRNA_arom conditions in female or male P0 cortex. This revealed that shRNA-arom reduced endogenous aromatase expression by ∼ 50% in both sexes. Data are represented as a percentage of control condition. Conditions were compared by two-way ANOVA; F(1,18)=95.59, p<0.0001, Bonferroni Post Hoc, ***, p < 0.001; n=8-12 sections from 4-6 pups/sex. Each data point represents an individual section. Scale bars = 20 μm.

Next, we assessed whether expression of shRNA_arom in the developing cortex by *in utero* electroporation resulted in a significant knockdown of aromatase in males and females. Embryos were electroporated with shRNA_arom expression construct or control shRNA (shRNA_scram) at E14.5 (**Figure 3 C and D**). Corrected integrated intensity measurements from confocal images taken of the P0 mouse cortex confirmed that shRNA_arom effectively reduced aromatase expression by approximately 50% compared to control condition in both sexes (**Figure 3C-E**). These data confirm the efficacy of aromatase knockdown by shRNA *in vivo*.

### Aromatase knockdown affects cortical migration divergently in male and females

To determine whether brain-synthesised estrogens play a role in the migration of neocortical cells, we assessed distribution of GFP+ cells in control and aromatase knockdown conditions (**Figure 4**). In P0 male mice, knockdown of aromatase resulted in an increase of GFP+ cells within the upper most portion of the CP with a concurrent reduction of GFP+ cells within the SVZ/VZ (**Figure 4 A and B**). Conversely, the opposite distribution was observed in the shRNA_arom condition in females; a decreased number of GFP+ was observed in the CP, where as an increase was detected in the SVZ/VZ (**Figure 4C and D**). Taken together, these data indicate that that knockdown of aromatase may accelerate radial neuronal migration in male, whereas migration is impaired in the female developing cortex.

**Figure 4.**
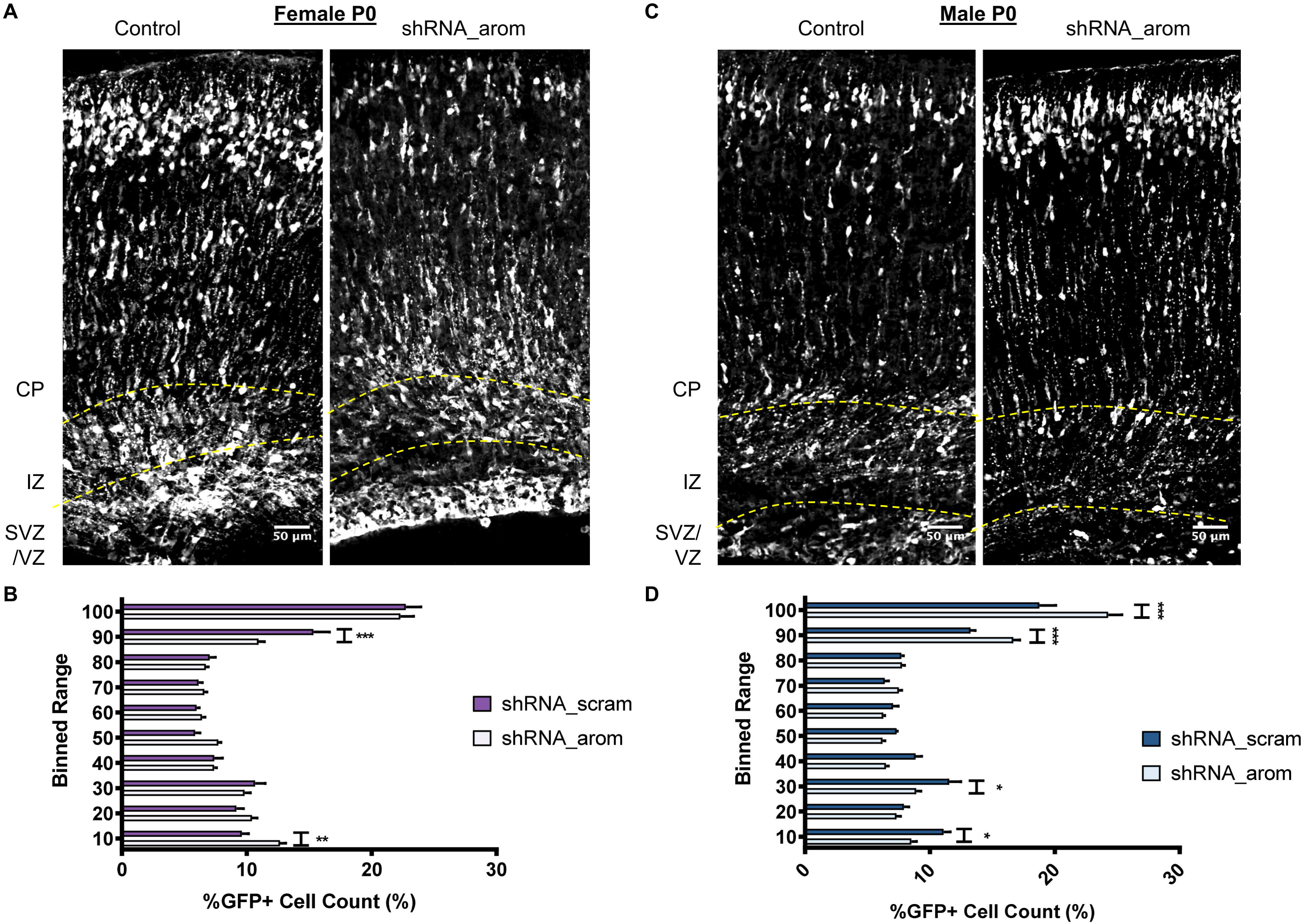
Loss of aromatase in developing cortex has divergent effects on neuronal migration in female and male cortex. **(A)** Representative confocal images of female P0 cortex *in utero* electroporated at E14.5 with either control (shRNA_scram) or shRNA_arom and stained for GFP. **(B)** Quantification of GFP+ cells distribution throughout the developing female cortex. In conditions where *Cyp19a1* had been knockdown (shRNA_arom), a significant more GFP+ cells were observed in the lowest bin, equating to the SVZ/VZ of the P0 cortex. Concurrently, a decrease in the number of GFP+ cells was measured in the upper most bins, equating to the upper portion of the CP. Conditions were compared by two-way ANOVA; F(9,140)=5.873, p<0.0001, Bonferroni Post Hoc, **, p<0.01, ***, p < 0.001; n=10-12 sections from 6-8 pups/condition/sex. **(C)** Representative confocal images of male P0 cortex *in utero* electroporated at E14.5 with either control (shRNA_scram) or shRNA_arom and stained for GFP. **(D)** Quantification of GFP+ cells distribution throughout the developing male cortex. Loss of aromatase induced by *Cyp19a1* knockdown (shRNA_arom) resulted in a significant increase of GFP+ cells in the upper portion of the CP. Congruent with this, a decrease in GFP+ cells was observed in the SVZ/VZ of the P0 male cortex. Conditions were compared by two-way ANOVA; F(9,160)=10.08, p<0.0001, Bonferroni Post Hoc, *, p<0.05, ***, p < 0.001; n=7-11 sections from 5-7 pups/condition/sex. Scale bars = 50 μm.

## Discussion

Aromatase is expressed in specific brain regions, where it controls the bioavailability of brain-synthesized estradiol within female and male brains (Saldanha *et al.*, 2011; Srivastava *et al.*, 2013; Lu *et al.*, 2019). Moreover, estradiol is present at significant levels in the brain of both sexes, including the developing cortex (MacLusky *et al.*, 1986; MacLusky *et al.*, 1994; Konkle & McCarthy, 2011). However, the functions of brain-synthesized estradiol during early corticogenesis development are unclear (Denley *et al.*, 2018). Here, we demonstrated that aromatase is widely expressed within the developing cortex of perinatal female and male mice. Aromatase expression was higher in males compared with females across the different laminae of the developing cortex. Interestingly, there were no differences in the pattern of migration in female and male cortices at P0. We used a knockdown approach to suppress endogenous *Cyp19a1* and thus aromatase expression in a subset of cortical progenitor cells destined to migrate to layer 2/3. Knockdown of aromatase had opposing effects on the migration of cortical progenitor cells in female and male developing brains. Specifically, loss of *aromatase* resulted in an increase of GFP+ cells in the CP with a concurrent decrease in the SVZ/VZ, indicating a potential increase in radial migration. Conversely, knockdown of aromatase in female mice resulted in a significantly decreased number of GFP+ cells in the upper layers of the developing cortex and an accumulation of cells within the SVZ/VZ, indicating a potential decrease in neuronal migration. Taken together, our data indicate that whilst there is no obvious difference in migration between male and females under control conditions, the influence of brain-synthesized estradiol on radial migration is sexually divergent and potentially involves the regulation of distinct mechanism in either sex.

The data presented in this study are consistent with a purported role for brain-synthesized estradiol in the development of the mammalian forebrain. Previous studies have demonstrated that high levels of aromatase are expressed in multiple regions of the brain, including the cortex (Beyer *et al.*, 1994; Yague *et al.*, 2008; Cisternas *et al.*, 2015). Importantly, the current results are consistent with this, but they further highlight that that there is greater expression of aromatase in the male developing cortex compared with females at the protein level. Although previous work implicated ERβ knockdown in the development of the cortex (Wang *et al.*, 2003), the current study provides evidence that brain-synthesised estradiol regulates neuronal migration in the developing cortex of both female and male mice. Whether systemic estradiol also impacts neuronal migration is unclear from these studies and would need to be studied further in the future.

A striking finding of this study is that the effect of *aromatase* knockdown on neuronal migration in the developing cortex is opposing in female and male mice. There are two possible explanations for this observation. First, estradiol could be exerting multiple effects on progenitor cells, such as controlling cell proliferation and/or apoptosis, as reported previously in the hypothalamus (Denley *et al.*, 2018; McCarthy *et al.*, 2018). Second, estradiol could be modulating the migration of newly born neurons in both sexes. No differences in the number of GFP+ cells were found under different conditions and between sex (data not shown), which indicates that the changes observed were dues to different localization of the labelled cells rather than increases or decreases in the overall number of cells. Thus, these effects are more likely an effect on the migration (or lack thereof) of newly born neurons. The small GTPase Rap1 mediates the migration, polarity, and establishment of neuronal morphology (Jossin & Cooper, 2011; Srivastava *et al.*, 2012b). Since we previously demonstrated that estradiol controls Rap1 activity in maturing cortical neurons (Srivastava *et al.*, 2008), it is possible that brain-synthesized estradiol mediates migration via a Rap1-dependent mechanism. It is also important to note that although knockdown approach using *in utero* electroporation allows us to examine the cell-autonomous effect of aromatase in the specific time critical for radial neuronal migration. However, we should be cautious in data interpretation, as off-target effect of shRNA may be confounded in the results. Thus, it is important to investigate the effect of aromatase(*Cyp19a1*)-haploinsufficiency in mouse models to elucidate aromatase-mediated mechanisms underpinning developmental phenotypes such as neuronal migration in the future.

The development of the cortex is fundamental for normal brain function. Interestingly, multiple animal models for autism spectrum disorders aimed at understanding the contribution of genetic and/or environmental risk factors to the underlying pathophysiology of this disorder have revealed that disruptions in early brain development is prevalent in these models. In particular, abnormalities in the development, migration, and organization of the developing cortex have been reported (Fenlon *et al.*, 2015; Varghese *et al.*, 2017). Moreover, there is accumulating evidence that elevated levels of fetal steroids, especially testosterone (Baron-Cohen *et al.*, 2015; McCarthy & Wright, 2017), are linked with autism spectrum disorders. Furthermore, rare mutations in the *CYP191A* gene have been reported in autistic patients and reduced ERβ and aromatase expression has been measured in autistic post-mortem tissue (Chakrabarti *et al.*, 2009; Sarachana *et al.*, 2011; Crider *et al.*, 2014). These lines of evidence have led to suggestions that altered steroidogenic activity and/or elevated levels of fetal testosterone could contribute to the pathophysiology of autism. It should be noted that knocking down *Cyp19a1* will both reduce estradiol levels and likely increase the levels of testosterone and other androgens within the developing cortex of these animals. Therefore, the current study may not only provide an insight into how reduced brain-synthesised estradiol levels impact development of the cortex, but also the impact of elevated levels of fetal testosterone on corticogenesisand therefore how dysregulation of fetal steroids could contribute to the emergence of neurodevelopmental disorders such as autism spectrum disorders.

In conclusion, the current study revealed that aromatase is expressed in the developing cortex of both female and male mice, and at higher levels in males than females. Knockdown of aromatase in cortical progenitor cells destined to migrate to layer 2/3, had marked sex-specific effects. Future studies focusing on understanding the mechanism underlying these effects, including investigating the potential role of Rap1, and also identifying the receptors that are responsible for the actions of the brain-synthesized estrogens (i.e., do brain-synthesized estradiols function via the classical “genomic” mode of action or do they act via a “membrane initiated” mode of action) are required. Together with the current work, these studies will help reveal the potential role of fetal steroids in normal development and how perturbations in this system may contribute to the emergence of disease.

## Conflict of Interest

The authors declare that the research was conducted in the absence of any commercial or financial relationships that could be construed as a potential conflict of interest.

## Author Contributions

K.J.S., A.S., A.K. and D.P.S. designed experiments. K.J.S. M.C.S.D., A.S. and D.P.S. performed all experiments and subsequent analysis. D.P.S. oversaw the study; K.J.S. M.C.S.D., and D.P.S. wrote the manuscript; all authors edited manuscript drafts.

## Funding

This work was supported by grants from Medical Research Council, MR/L021064/1, Royal Society UK (Grant RG130856), the Brain and Behavior Foundation (formally National Alliance for Research on Schizophrenia and Depression (NARSAD); Grant No. 25957), awarded to D.P.S.; NIH grants R01DA041208 (A.K.), P50MH094268 (A.K.), Catalyst award (A.K.), and Brain & Behavior Research Foundation (A.K., A.S.); K.J.S. was supported by a McGregor Fellowship from the Psychiatric Research Trust (Grant McGregor 97) awarded to D.P.S.; K.J.S. is the recipient of an Institute of Psychiatry, Psychology and Neuroscience, Independent Researcher Early Career Award.

